# SIRT2-knockdown Rescues GARS-induced Charcot-Marie-Tooth Neuropathy

**DOI:** 10.1101/854588

**Authors:** Chao Shen, Qian Qi, Yicai Qin, Dejian Zhou, Xinyuan Chen, Yun Qi, Zhiqiang Yan, Xinhua Lin, Jinzhong Lin, Wei Yu

## Abstract

Charcot-Marie-Tooth disease is the most common inherited peripheral neuropathy. Dominant mutations in glycyl-tRNA synthetase (GARS) gene cause peripheral nerve degeneration and lead to CMT disease type 2D. Mutations in GARS (GARS^CMT2D^) show partial loss-of-function features, suggesting that tRNA-charging deficits play a role in disease pathogenesis, but the underlying mechanisms are not fully understood. In this study we report that wild-type GARS tightly binds the NAD^+^-dependent deacetylase SIRT2 and inhibits its deacetylation activity, resulting in the hyperacetylated α-tubulin, the major substrate of SIRT2. Previous studies showed that acetylation of α-tubulin protects microtubules from mechanical breakage and keep axonal transportation. However, CMT2D mutations in GARS can not inhibit SIRT2 deacetylation, which leads to decrease acetylated α-tubulin and severe axonal transport deficits. Genetic reduction of *SIRT2* in the *Drosophila* model rescues the GARS–induced axonal CMT neuropathy and extends the life span. Our findings demonstrate the pathogenic role of SIRT2-dependent α-tubulin deacetylation in mutant GARS-induced neuropathies and provide new perspectives for targeting SIRT2 as a potential therapy against hereditary axonopathies.

## Introduction

Charcot-Marie-Tooth disease is the most common inherited peripheral neuromuscular disorder, which affects 1 of every 2500 persons[1, 2]. A well-organized microtubule network is required for peripheral axons to effectively transport vesicles between the soma and the synapse. The acetylation of α-tubulin within the microtubules promotes the anchoring of the molecular motor kinesin and stimulates vesicular transport along the microtubules. Additionally, acetylated microtubules are far more stable and resistant to drug-induced depolymerization than non-acetylated microtubules[3]. Defects in axonal transport are often associated with neurodegeneration and with peripheral neuropathies in particular[4].

SIRT2 belongs to the class III HDACs and controls the acetylation status of several proteins, including α-tubulin[5]. Targeting the activity of SIRT2 has been shown to be beneficial in Parkinson’s disease (HD) and Huntington’s disease (HD) but not in CMT[6]. Although SIRT2 is able to deacetylate α-tubulin *in vitro*[5], knockout of SIRT2 in mice does not alter the acetylation levels of α-tubulin *in vivo*[7, 8], indicating that SIRT2 may deacetylate α-tubulin under special conditions. For example, SIRT2, not HDAC6, is responsible for deacetylation of α-tubulin during inflammasome activation in mice macrophages[9]. Here we provide another evidence of SIRT2 knockdown rescuing the GARS-induced CMT diseases.

Dominant mutations in the glycyl-tRNA synthetase (GARS) gene cause peripheral nerve degeneration and lead to type 2D CMT disease. Notably, genetic studies in mice models of GARS^P234KY^ and GARS^C157R^ and a fly model of GARS^G240R^ revealed that these dominant mutations in GARS could cause CMT through toxic gain-of-function effects[10, 11]. Several studies have demonstrated that mutations in GARS (GARS^CMT2D^) that result in tRNA-charging deficits play a role in disease pathogenesis, suggesting that partial loss of aminoacylation activities are involved[12, 13], but the underlying mechanisms are not fully understood.

In this study, we report that wild-type GARS tightly binds the NAD^+^-dependent deacetylase SIRT2 and inhibits its deacetylation activity, resulting in hyperacetylated α-tubulin, which is the major substrate of SIRT2. Previous studies showed that the acetylation of α-tubulin protects microtubules from mechanical breakage[14, 15] and maintains axonal transportation[16]. However, CMT2D mutations in GARS alter its structural conformation, enabling GARS^CMT2D^ to lose binding with SIRT2 and reduce the inhibition of SIRT2 deacetylation, which leads to decrease acetylated α-tubulin and severe axonal transport deficits. The knockdown of *SIRT2* in a *Drosophila* model rescues GARS–induced axonal CMT neuropathy and extends the life span. Our findings demonstrate the pathogenic role of SIRT2-dependent α-tubulin deacetylation in mutant GARS-induced neuropathies and provide new possibilities for targeting SIRT2 as a potential therapy against hereditary axonopathies.

## Results

### Wild-type GARS tightly binds the SIRT2 to inhibit its activity, not GARS^CMT2D^

Since targeting SIRT2 has been shown to benefit in other neurodegeneration diseases[6]. We want to test whether SIRT2, a major deacetylase of α-tubulin, might also play the critical role in CMT diseases. We performed a co-immunoprecipitation assay to determine whether SIRT2 might interact with GARS or GARS^CMT2D^ mutants. First, we transiently cotransfected HEK293T cells with FLAG epitope-tagged GARS (wild-type and CMT2D mutants) and SIRT2-HA. SIRT2 interacted with GARS^WT^ but not with the GARS^CMT2D^ mutants (G526R, E71G) (Figure 1A). In a similar reverse analysis, SIRT-FLAG could pull down wild-type GARS-Myc but not the GARS^CMT2D^ mutants (Figure 1B). These results indicated that only GARS^WT^, not GARS^CMT2D^ could interact with SIRT2. To determine whether the interaction of GARS^WT^ with SIRT2 would directly affect the deacetylase activity of SIRT2, we purified SIRT2 and GARS from *E. coli* to perform an *in vitro* deacetylase assay. Using seryl-tRNA synthetase (SARS), which is known to bind to SIRT2 and promote its deacetylase activity[17], as a control, we tested the effect of different GARS concentrations on SIRT2 activity (Figure 1C). As expected, SARS increased the SIRT2 activity at a ratio 1:1, while GARS decreased SIRT2 activity at a ration 2:1 (GARS:SIRT2). Increasing the GARS:SIRT2 ratio to 4:1 and 6:1 did result in additional reduction. While the same concentrations of GARS^CMT2D^ barely inhibited SIRT2 deacetylase activity (Figure 1D). The data demonstrate that the GARS interaction directly inhibits SIRT2 deacetylase activity in a dose-dependent manner.

**Figure 1.**
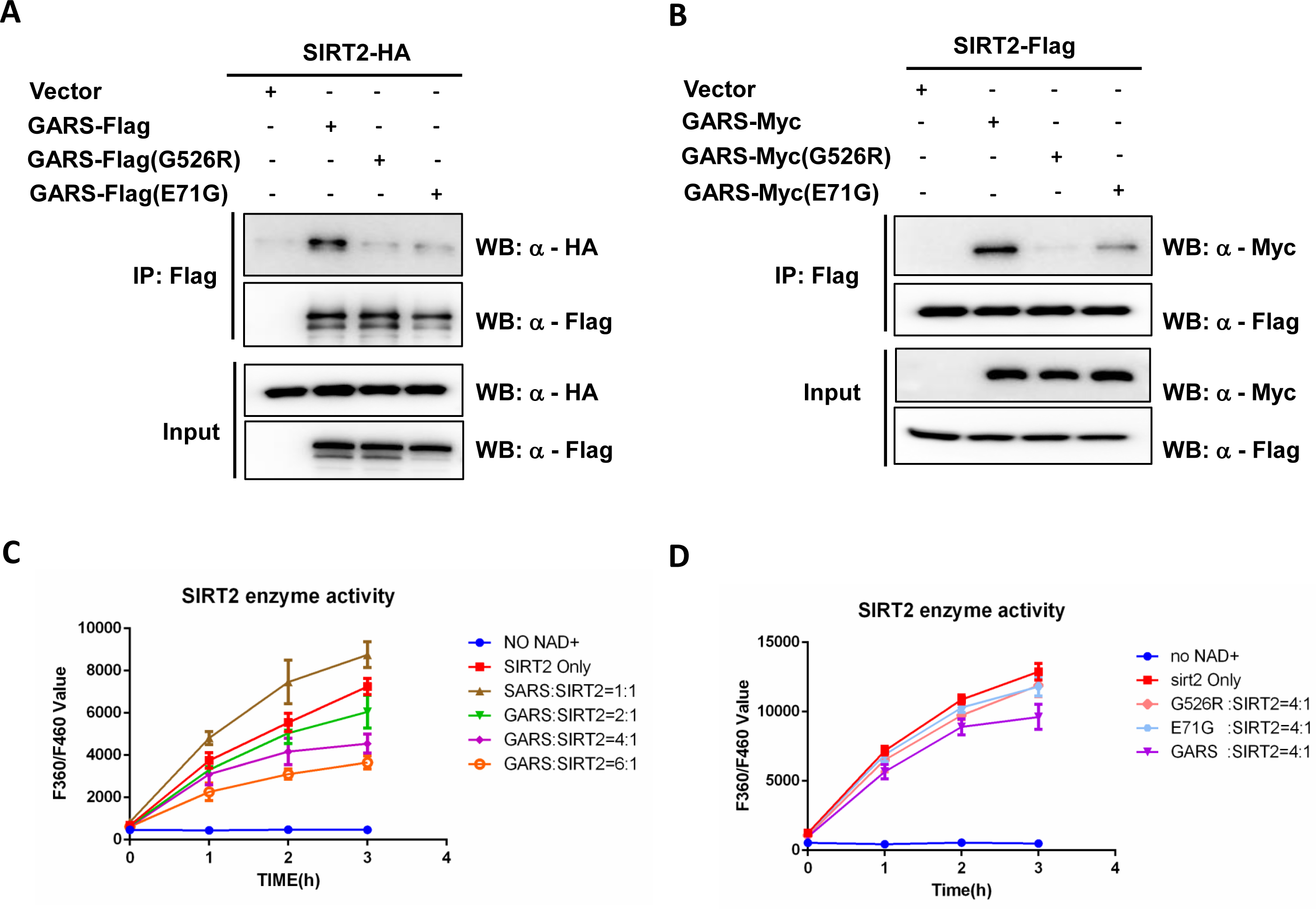
Wild-type GlyRS tightly binds the SIRT2 to inhibit its activity, not GlyRS^CMT2D^. (A-B) SIRT2 interacts with wild-type GlyRS, not GlyRS mutant. SIRT2 or GlyRS were immunoprecipitated from HEK293T cell lysates with Flag beads. Precipitated GlyRS-Flag or GlyRS mutant (G526R, E71G) was detected by anti-Flag antibody and co-IP SIRT2-HA was detected by anti-HA as indicated(A). Precipitated SIRT2-Flag was detected by anti-Flag antibody and co-IP GlyRS-Myc or GlyRS mutant (G526R, E71G) was detected by anti-Myc as indicated (B). (C-D) Effect of GlyRS(C) or GlyRS mutant (G526R, E71G) (D) on SIRT2 deacetylation activity. SIRT2 (1 µM) were incubated with purified GlyRS or GlyRS (G526R, E71G) (concentration measured as monomer) at the indicated ratios. The deacetylase activities of recombinant human SIRT2 were measured by monitoring the fluorescence intensity (excitation at 360 nm and emission at 460 nm) using a substrate peptide with one end coupled to a fluorophore and the other end to a quencher. SIRT2 incubated with SerRS was performed as a positive control(C). A reaction without NAD^+^ was performed as a negative control. Data are presented as mean±s.d., n=3 wells, from three independent experiments.

### Wild-type GARS inhibits SIRT2 deacetylation activity on acetylated α-tubulin, not in GlyRS^G526R^ mutants

To understand whether the knockdown of GARS might affect deacetylation of SIRT2 on α-tubulin, we detected the level of α-tubulin acetylation in HEK293T cells expressing two different GARS siRNA. The acetylation levels of α-tubulin in the two different GARS knockdown cells were significantly decreased and were similar to those resulting from the overexpression of SIRT2 in HEK293T cells (Figure 2A). To demonstrate that the inhibition of SIRT2 affects the acetylation of α-tubulin, we performed an *in vitro* assay with total cellular lysates with increased concentrations of wild-type GARS and GARS^G526R^. We observed a greatly increased levels of α-tubulin when cells were incubated with wild-type GARS in a dose-dependent manner but not in the GARS^G526R^ (Figures 2B-2C). These data indicate that the interaction of wild-type GARS and SIRT2 inhibits the deacetylase activity of SIRT2 and leads to the hyperacetylation of α-tubulin, which might protect microtubules from mechanical breakage. To gain mechanistic insights into the role of CMT2D mutation in GARS, we inspected the 3D structure of the human GARS protein (PDB: 2ZT5) and found both CMT2D mutations caused a conformational opening surface in GARS (Figure 2D). Comparing with E71 in WT having a negatively charged regions, E71G mutant represents neutral charged regions (Figure 2E), which made us to hypothesize that structural alteration induced by CMT2D mutations in GARS might disrupt the proper binding with SIRT2.

**Figure 2.**
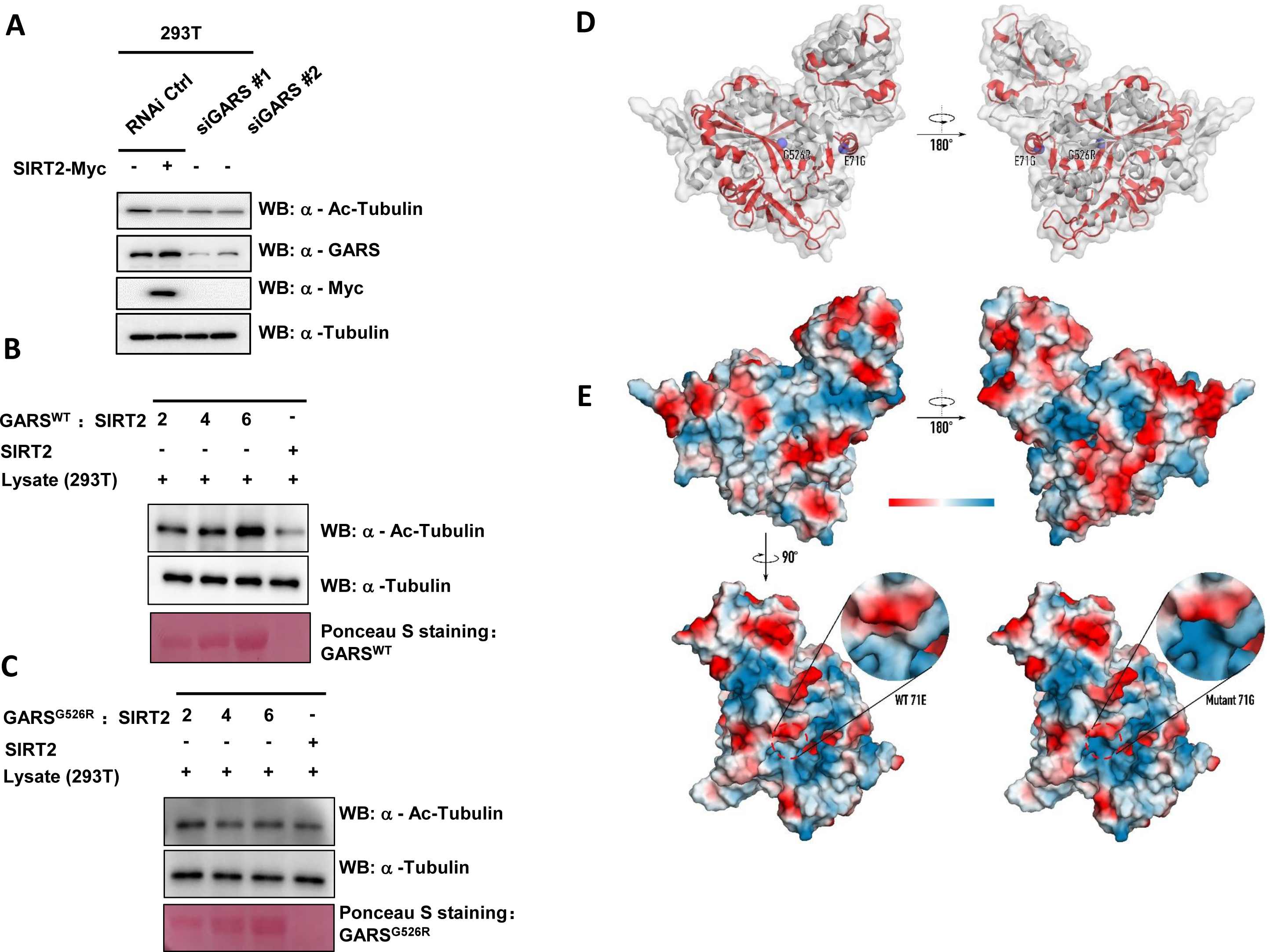
Wild-type GARS inhibits SIRT2 deacetylation activity on acetylated a-tubulin, not in GlyRS^G526R^ mutants. (A) Both knockdown of GlyRS and overexpression of SIRT2 result in decreased tubulin acetylation. HEK293T cells were transfected with GARS siRNA or control siRNA 24h later, SIRT2-Myc was overexpressed in cells transfected with control siRNA. After an additional 48h, Cells were harvested and acetylation of tubulin was detected by western blot. (B-C)The recombinant SIRT2 and recombinant GlyRS^**WT**^ (B) or recombinant GlyRS^**G526R**^(C) was incubated with cellular lysate derived from 293T cells *in vitro*. The reaction products were detected by Western blotting for acetylated tubulin, α-tubulin, and His-tag. 1/2, 1/4, 1/6: The total amount of GlyRS is twice, four times, six times that of SIRT2 respectively. (D) Structure of human GARS (PDB: 2ZT5) with opened surface (red) caused by G526R and E71G mutations. (E) Electrostatic potential maps of WT GARS or GARS with E71G mutation. All images generated with PyMol.

### SIRT2 knockdown rescues CMT phenotype in GlyRS^G526R^ flies

A previous study showed that GARS with a CMT-associated mutation induced motor performance deficits in a *Drosophila* model[18]. To determine how SIRT2 affects a GARS mutant *in vivo*, we next assessed the motor behavior resulting from *sirt2*-knockdown in GARS^G526R^ flies. We generated SIRT2 knockdown in GARS^G526R^ flies (Extend Figure 1A) and confirmed the knockdown efficiency of SIRT2 by realtime-PCR and western blot (Extend Figure 1B-1C). Compared to untreated GARS^G526R^ flies, GARS^G526R^ flies with *sirt2*-knockdown showed significantly restored climbing ability and similar to WT flies (Figure 3A, extend Figure 2A). To provide further evidence that the inhibition of SIRT2 might rescue the phenotype of CMT-mutant GARS in the *Drosophila* model, we fed GARS^G526R^ flies with AGK2, which is a specific inhibitor of SIRT2, and then examined the climbing ability of these treated GARS^G526R^ flies. We found that 100 μM AGK2-fed GARS^G526R^ flies showed significantly shorter climbing times compared with DMSO-fed GARS^G526R^ flies (Extend Figure 2B). Interestingly, the motor performance of GARS^G526R^ flies was gradually rescued by AGK2 treatment in GARS^G526R^ flies, in a time-dependent manner (Figure 3B). Since mutant GARS flies also showed neuronal morphological defects, we performed a staining analysis of neuromuscular junctions (NMJs) to assess the development status of NMJs in *sirt2*-knockdown GARS^G526R^ flies. Consistent with a previous report that GARS^G526R^ flies showed a dramatic decrease in NMJs, we found that *sirt2*-knockdown in GARS^G526R^ flies significantly increased the numbers of NMJs in third instar larvae to levels similar to those in WT flies (Figures 3C-3D). Moreover, we observed the acetylated tubulin was significantly rescued in *sirt2*-knockdown in GARS^G526R^ flies comparing with the GARS^G526R^ flies (Figure 3E). Overall, these results strongly indicate that the inhibition of SIRT2 could rescue the motor performance deficits in GARS-mutant CMT flies in a *Drosophila* model.

**Figure 3.**
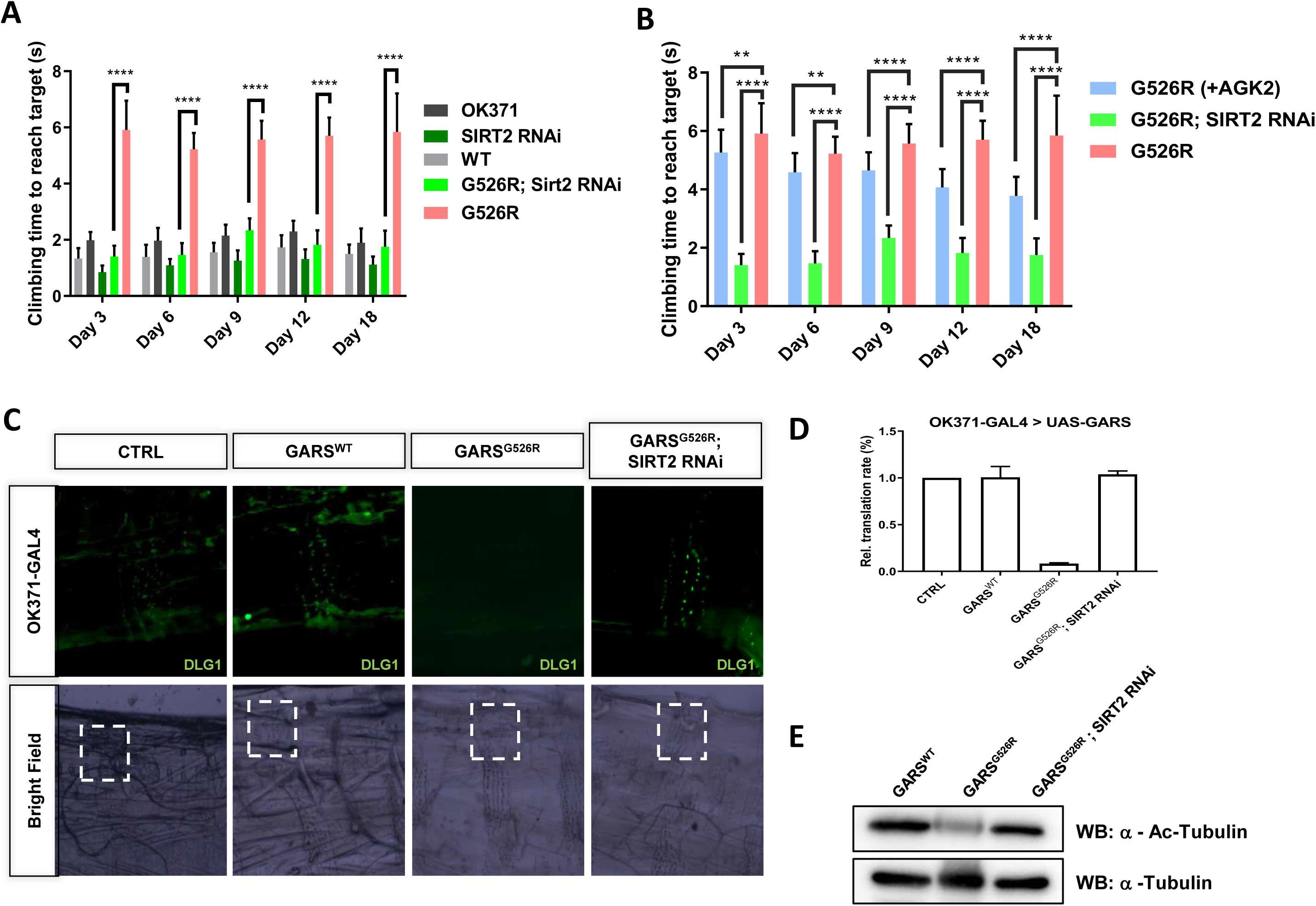
SIRT2 knockdown rescues CMT phenotype in GlyRS^G526R^ flies. (A) Bar graph displaying average climbing time to reach target in a negative geotaxis assay of female flies in motor neurons (OK371-GAL4). WT (gray), SIRT2 RNAi in GARS^G526R^ (green) and GARS^G526R^ (pink). N>100. Error bars represent s.e.m. ****P<0.0001. (B) 100 M AGK2 was fed from 12h flies to 3-to 18-day-old flies, then motor performance was detected and bar graph was displayed average climbing time. AGK2-fed GARSG526R (blue), SIRT2 RNAi in GARSG526R (green) and GARSG526R (pink). 4% DMSO was fed in all lines. N>100. Error bars represent s.e.m. ****P<0.0001. (C) SIRT2 knockdown restore NMJ in GlyRS^**G526R**^ mutants. NMJs of third instar larvae expressing GARS in motor neurons (OK371-GAL4) were visualized by staining for the postsynaptic marker discs large 1 (dlg1). Results indicate the NMJ on muscle 4, which is missing in GARS^**G526R**^ flies and is appearing in SIRT2 knockdown GARS^**G526R**^ flies. (D) Quantification of the percentage of animals with muscle 4 innervated. N=3. Error bars represent s.e.m. (E) Western blot analysis of SIRT2 in lysate of different fly lines.

### SIRT2 knockdown restores life span in GlyRS^G526R^ flies

A previous study showed that CMT-mutant GARS flies have shortened lifespans[18]. To investigate whether SIRT2 knockdown or inhibition by AGK2 could affect the lifespan of CMT-mutant GARS flies, we conducted a longevity analysis of *sirt2*-knockdown GARS^G526R^ flies using a ubiquitously expressed driver (Tub > SIRT2^RNAi^) or 200 μM AGK2-fed GARS^G526R^ flies. We first generated ubiquitous GARS transgene expression flies using the GAL80 target system. Consistent with a previous report that GARS^WT^ files did not show changes in their lifespan, but GARS^G526R^ flies showed significantly reduced lifespans compared to GARS^WT^ files. Strikingly, the knockdown of *sirt2* expression in GARS^G526R^ flies extended the lifespans which was similar to that observed in GARS^WT^ files (Figure 4A). Also AGK2-fed GARS^G526R^ flies also showed significantly extended lifespans compared to GARS^G526R^ files (Figure 4A). These data indicate that the loss of SIRT2 function in CMT-mutant GARS flies resulted in a longer lifespan and strongly indicated the involvement of SIRT2 in CMT phenotype regulation.

**Figure 4.**
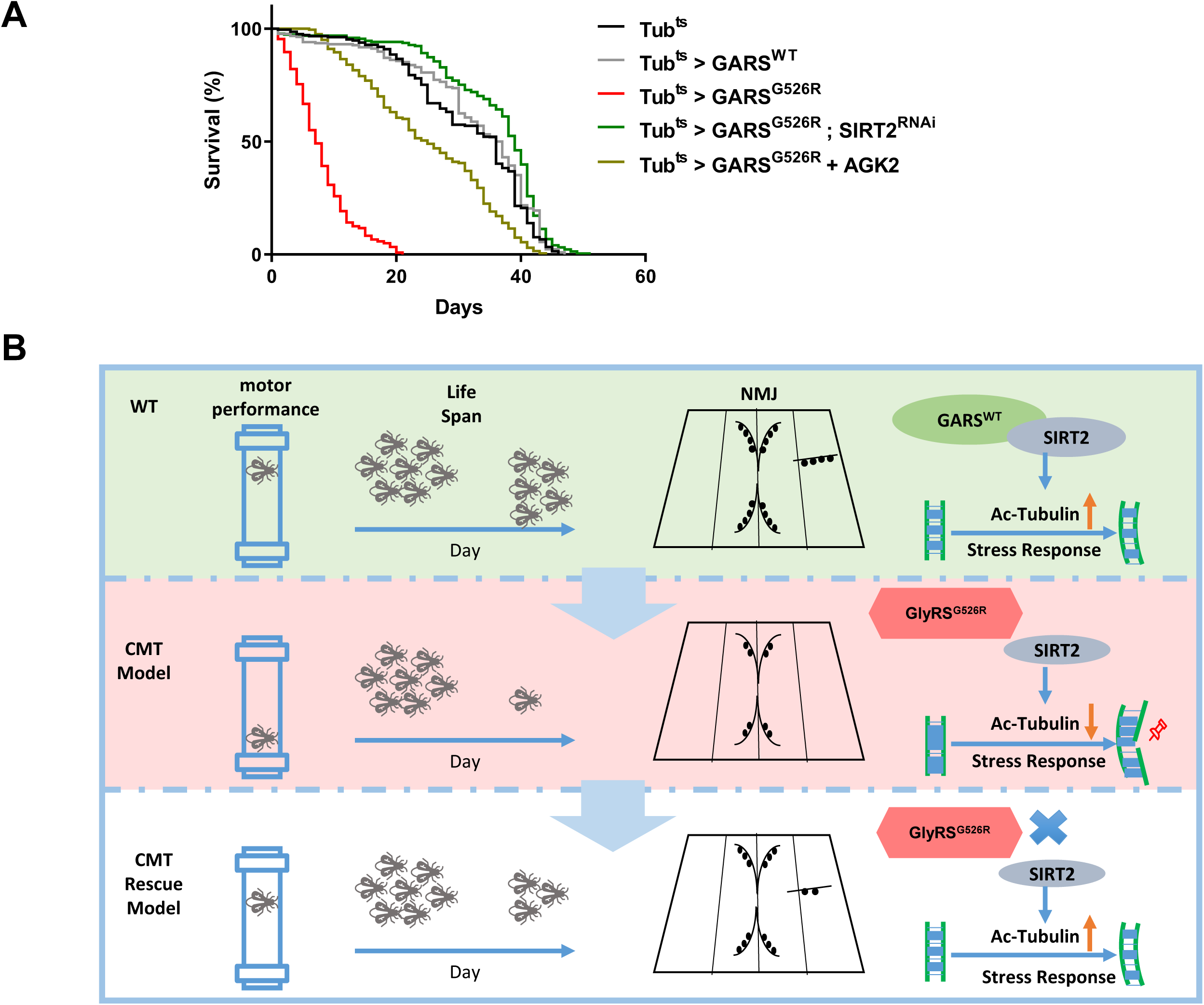
SIRT2 knockdown restore life span in GlyRS^G526R^ flies. (A) Kaplan–Meier survival curves displaying the lifespan of male flies from the adult stage onwards. N > 200. (B) Working model

## Disscussion

Targeting the expression or deacetylation activity of SIRT2 has been shown to benefit in several neurodegenerative diseases such as Parkinson’s disease, Huntington’s disease and amyotrophic lateral sclerosis (ALS), but not shown benefit in CMT[6]. Mutations in five aminoacyl tRNA synthetases, including GARS, have been identified that cause CMT or related peripheral neuropathies[19]. In this study, we showed that WT GARS but not GARS^CMT2D^ binds to SIRT2 and prevents its deacetylation activity, which leading to the hyperacetylation of α-tubulin. Our data revealed that CMT2D mutations in GARS can not inhibit the deacetylation activity of SIRT2, resulting in hypoacetylated α-tubulin and severe axonal transport deficits. Furthermore, the genetic reduction of *SIRT2* expression and a specific inhibitor of SIRT2 in a *Drosophila* model rescued the GARS–induced axonal CMT neuropathy and extended the life span (Figure 4B). Our study outlines a method for targeting SIRT2 to reduce the GARS-induced CMT neuropathy.

Acetylated α-tubulin has been shown to play multiple cellular functions, including the regulation of cell motility and polarity and participation in intracellular transport, ciliary assembly and immune and viral responses[20]. Interestingly, the Maxence Nachury group recently reported that acetylated α-tubulin protects microtubules from mechanical breakage[14, 15], which may be linked to the neuropathy of CMT disease, which usually occurs in the first two decades of life because this stage in life, the hypoacetylated microtubules in peripheral motor neurons are not able to endure the mechanical stress and accumulated the lattice damage, resulting in the degeneration of peripheral motor and sensory axons. Hence, we demonstrated that SIRT2 knockdown could ameliorate the phenotype associated with CMT2D pathology by rescuing neuromuscular junctions in flies. This is an interesting outcome, since defects in neuromuscular junction maturation were shown to precede the impaired connectivity of lower motor neurons in CMT2D mice[21].

SIRT2 localizes in the cytoplasm and was identified as the deacetylase of lysine 40 in α-tubulin *in vitro*[5]. Compared with HDAC6, SIRT2 may function as a deacetylase of α-tubulin in special conditions, such as in the mitotic spindle[22] or during the inflammasome activation in macrophages[9]. The inhibition of SIRT2 expression or deacetylase activity was shown to rescue a-synuclein toxicity in a model of Parkinson’s disease[23]. Additionally, the pharmacological inhibition of SIRT2 could impair sterol biosynthesis to provide neuroprotection in models of Huntington’s disease[24]. Our study provides another promising strategy to treat the peripheral neuropathy resulting from CMT by inhibiting SIRT2, suggesting that the targeting of SIRT2 might have a general beneficial impact on neurodegenerative diseases.

Currently, two genetics experiments in flies and mice have demonstrated that dominant mutations in GARS cause CMT through toxic gain-of-function effects[10, 18]. Several binding partner of CMT mutants have been discovered, including Nrp1[25], Trk[26] and HDAC6[27, 28]. Here we presented that SIRT2, the acetylated tubulin deacetylase, is able to bind tightly with wildtype GARS comparing with CMT mutants, we didn’t rule out the possibility that CMT mutants might gain the functions to increase deacetylation activity of SIRT2 which leading to decrease the acetylated tubulin. Interestingly, HDAC6 is able to form a complex with SIRT2[5] which made us to hypothesize that CMT mutants might bind with HDAC6 to affect deacetylation activity of SIRT2. This definitely needs to be addressed in future experiments.

## Materials and Methods

### Antibodies

Anti-SIRT2 antibody was purchased from Abcam (ab51023). Anti-GlyRS antibody was purchased from Proteintech (15831-1-AP). Antibodies against α-tubulin and acetylated tubulin at Lys40 (TubulinK40Ac) were purchased from Invitrogen (62204) and Sigma (T6793), respectively. Antibodies to FLAG (Sigma, SAB4301135), HA (Abcam, ab9110), Myc (Abcam, ab9106) and His (abmart 293670) were commercially obtained.

### Plasmid Construction

SIRT2 were cloned to pCDNA3.1-FLAG vector. Point mutations for GlyRS were generated by site-directed mutagenesis (Toyobo KOD Mut Kit). Human GlyRS and SIRT2 genes were cloned into the pET-28a(+) to express with a C-terminal his-tag in Escherichia coli.

### Cell Culture and Transfection

HEK293T (human embryonic kidney) cells were grown in DMEM medium supplemented with 10% (vol/vol) 10% fetal bovine serum. Cultures were maintained at 37 °C in a humidified atmosphere containing 5% (vol/vol) CO2. Cells were grown to 50 to 70% confluency before treatments. Cell transfection except for siRNA was carried out by polyethylenimine (PEI) according to the manufacturer’s protocol. Cell transfection for siRNA was carried out by Lipofectamine 2000 according to the manufacturer’s protocol.

The sequences of siRNAs were as follows:

siRNA_GARS1, 5′-CTTGAGACCAGAAACTGCA-3′;

siRNA_GARS2, 5′-GTAGCTGAGAAACCTCTGA-3′;

### Coimmunoprecipitation assays

SIRT2 and GlyRS or mutant GlyRS were co-transfected into HEK293T cells, and cell lysates were immunoprecipitated with Flag beads (Sigma Aldrich) overnight at 4 °C, and then boiled with SDS loading buffer and subjected to Western blotting. SIRT2-Flag and MYC-tagged GlyRS or mutant GlyRS were detected as indicated.

### Cell Lysis and Western Blot Analysis

For cell-based experiments, cells were washed three times in PBS and resuspended with lysis buffer containing 50 mM Tris, pH 7.5, 150 mM NaCl, 1% Nonidet P-40, 1 mg/mL aprotinin, 1 mg/mL leupeptin, 1 mg/mL pepstatin, 1 mM Na_3_VO_4_, and 1 mM phenylmethylsulfonylfluoride (PMSF),25 mM NAM and trichostatin A (TSA) for 20 min and centrifuged for 20 min at 12,000 rpm; the insoluble fraction was discarded. The lysates were fractionated by SDS/PAGE and transferred to nitrocellulose filter (NC) membranes. The membranes were blocked for 1 h with Tris-buffered saline with Tween 20 (TBST) containing 5% (mass/vol) nonfat dry milk. After incubation with primary antibodies (anti-acetylated tubulin (K40) (Proteintech), anti-α-tubulin (Invitrogen), anti-SIRT2(Abcam) or anti-His (Abmart), each diluted 1:1000), the membranes were washed and incubated with HRP-conjugated anti-mouse (Cell Signaling Technology) or anti-rabbit secondary antibodies (Cell Signaling Technology), followed by detection using ECL Western blotting substrate (Bio-Rad).

### SIRT2 deacetylase activity assay

The reaction buffer contains 50 mM Tris-HCl (pH 9.0), 4 mM MgCl_2_, 0.5 mM DTT, 1µM MAL(Boc-Lys(AC)-AMC), 1 mM NAD+, 1µM recombinant SerRS, 1 µM recombinant SIRT2, and recombinant GlycRS or GlyRS mutant at different ratio indicated in Figure 1, 1b. The reactions were performed at room temperature. Add the stop solution (0.05 g/ml trypsin) of the same volume as the reaction system to stop the reaction, and measure fluorescence intensity as indicated.

### Purification of Recombinant SIRT2, GlyRS^WT^ and GlyRS^CMT2D^ mutants

All genes were PCR-amplified and cloned into the pET.28a vector to produce His6-tag-fused recombinant proteins. Point mutations were introduced by the site-directed mutagenesis approach. All recombinant proteins used in this study were expressed in E. coli BL21 (DE3) induced by 0.5 mM IPTG at 16°C for 20 hr and collected by sedimentation. To purify wild-type GlyRS and the G526R GlyRS mutants, the E. coli cells were resuspended in binding buffer (20 mM Tris-Cl pH 8.0, 500 mM NaCl and 25 mM imidazole), lysed with a high-pressure homogenizer and sedimented at 18000 rpm for 1 hr to pellet the debris. The supernatant lysates were purified by HisTrapTM FF. Then the proteins were further purified on an AKTA purifier (GE Healthcare), and eluted with elution buffer (20 mM Tris-Cl pH 8.0, 500 mM NaCl and 500 mM imidazole). All the purified proteins were concentrated by centrifugal filtrations, then stored in aliquots at -80°C.

### In Vitro α-Tubulin Deacetylation Assay

For SIRT2 experiments (Figure 2), the corresponding ratio (1/2, 1/4, 1/6) of recombinant SIRT2 (2.5μg) and GlyRS^WT^ (1.5μg/μl) or GlyRS^CMT2D^ mutants (1.8μg/μl) were resuspended in 100 μl of SIRT deacetylase buffer (50 mM Tris-HCI, 4 mM MgCl_2_, 0.2 mM DTT, [pH 9.0]) added with 100 μg of total cellular lysate from untransfected NSC-34 cells. Reactions were preincubated for 15 min and incubated for 1 hr at 37°C after addition of 1 mM NAD. A reaction is added with 5 mM nicotinamide as negative control. Reactions were stopped by adding 34 μl of 5× SDS-PAGE loading buffer. 8 μl of each reaction was separated on 12% SDS-PAGE gels and Western blotted as described above.

### Drosophila Genetics

The OK371-GAL4 driver line was kindly provided by Pro. Junhai Han from Southeast University and other GAL4 driver lines were obtained from Prof. Xinhua Lin from Fudan University. The dGARS deficiency lines described in this paper were kindly gifted from Prof. Erik Storkebaum in Max Planck institute for Molecular Biomedicine. SIRT2-RNAi lines were bought from Tsinghua Fly Center.

### Lifespan analysis

For determination of adult offspring frequencies, the number of adult flies eclosing was counted for each genotype. For assaying longevity, the tub-Gal4 driver was combined with a ubiquitously expressed temperature-sensitive Gal80 inhibitor (tub-Gal80ts). Fly crosses were cultured at 19°C and adult progeny carrying the tub-Gal4, tub-Gal80ts and UAS-GARS transgenes were shifted to 30°C to induce transgene expression. Males were collected within 24 h of eclosion and grouped into batches of 20 flies per food vial. The number of dead flies was counted every day and flies were transferred to fresh food vials every 2 days. At least 200 flies per genotype were used.

### Analysis of neuromuscular junction

For observation of neuromuscular junction, it was dissected from third instar larvae that selectively express target genes in motor neurons (OK371-GAL4). Larval filets were prepared and stained with primary antibodies against dlg1 (DSHB. 1/200).

### Drosophila behavior analysis

Flies for motor performance assays were kept at 25°C with a 12-h light/dark cycle. Female flies were collected within 24 h after eclosion and divided into groups of 20 individuals. Motor performance of 3-to 18-day-old flies was evaluated. On the day of the assay, flies were transferred in test tubes without anaesthesia and assayed within 15 min under standardized daylight conditions. Three test tubes were loaded into a self-made device, which was released from a certain height. The device fell down onto table, shaking the flies to the bottom of the test tubes and inducing a negative geotaxis climbing response. The whole procedure was videotaped with a camera and repeated five more times. Average climbing time to reach 6 cm was determined and compared between genotypes. At least 100 flies per genotype were used.

### Isolation of total RNA from fly larvae

Total RNA was isolated from third instar larvae under acidic conditions using Trizol. Approximately 10 frozen larvae were covered with 5 mL liquid nitrogen. All subsequent steps were carried out on ice or at 4°C. And 1 mL of Trizol were added to the homogenate, and the mixture was vortexed for 30 s followed by chilling on ice for 5 min, an amount of 0.3 mL of chloroform was added and mixing the samples repeatedly. After centrifugation at 12,000g for 10 min, the aqueous phase was transferred to a new tube. RNA was precipitated by 0.5 mL isopropanol by chilling on ice for 10 min. The resulting RNA pellet was washed twice with 75% ethanol. RNA was resuspended in DEPC and stored at 80°C.

### Statistical analysis

We used GraphPad Prism 6 software for statistical calculations. Unpaired t-test was used to analyze offspring frequency and behavior data. All data are reported as the mean SEM.

## Fly information

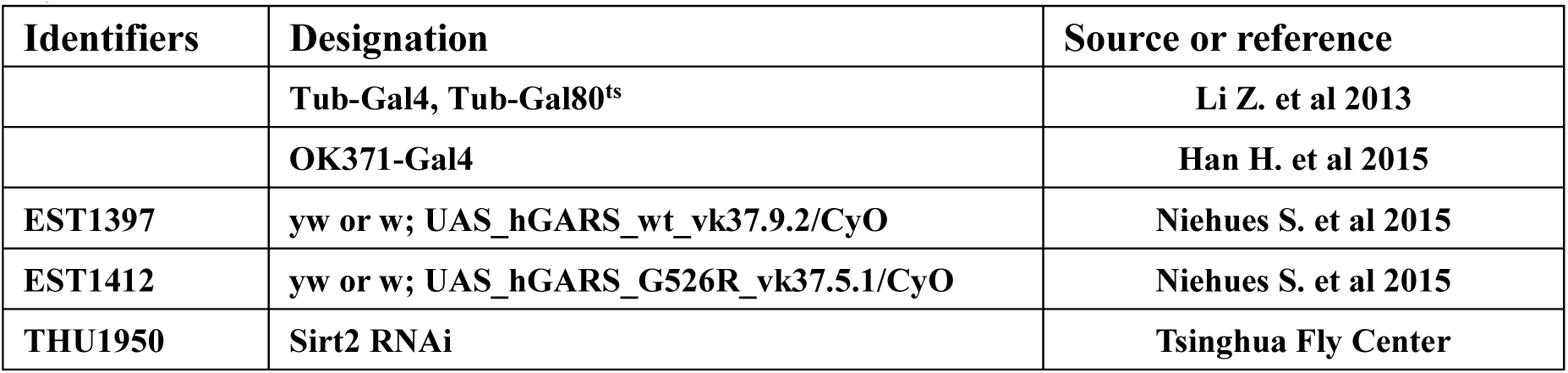

## Acknowledgements

We thank the members of Yu Lab for discussions and supports throughout this study. The dGARS deficiency lines described in this paper were kindly gifted from Prof. Erik Storkebaum in Max Planck institute for Molecular Biomedicine. The OK371-GAL4 driver line was kindly provided by Pro. Junhai Han from Southeast University. We thank the facility center of State Key Laboratory of Genetic Engineering for assistance on enzymatic activity experiments. This work was supported by the National Key Research and Development Program of China (2016YFA0500600), National Natural Science Foundation of China (31771545, 91749120). W.Y. was supported by China ‘‘Thousand Youth Talents’’ and the Program for Professor of Special Appointment (Eastern Scholar) at Shanghai Institutions of Higher Learning.

## Author contributions

### Conceptualization

Wei Yu.

### Biochemistry

Chao Shen, Qian Qi, Yicai Qin and Xinyuan Chen.

### Structural

Dejian Zhou and Jinzhong Lin.

### Fly experiments

Chao Shen, Qian Qi, Yun Qi, Ziqiang Yan, Xinhua Lin.

### Writing

Chao Shen, Qian Qi, Yicai Qin and Wei Yu wrote the paper.

---

**Supplemental Figure 1.**
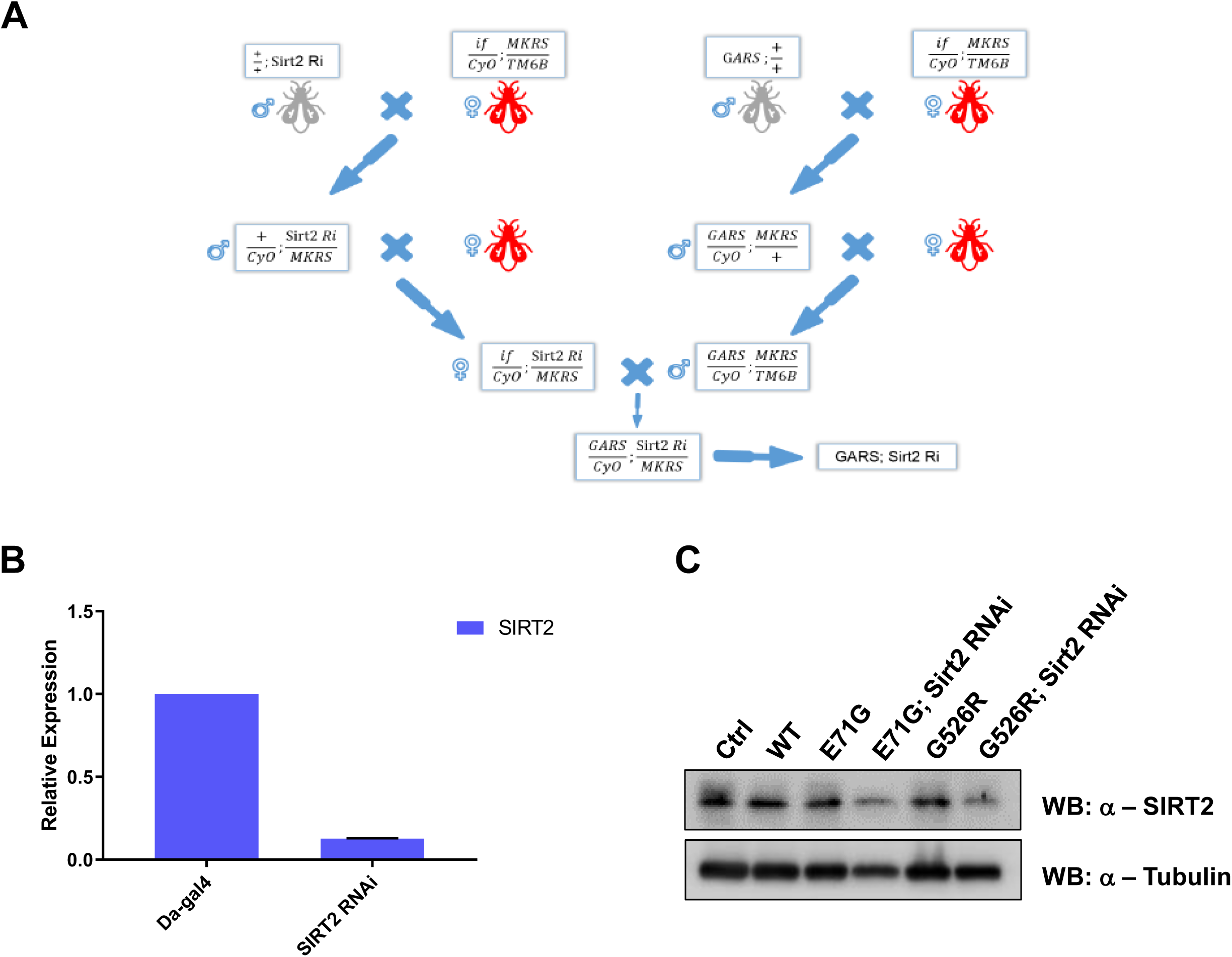
(A) The pipeline of construction of SIRT2 Knockdown in GARS^CMT^ flies. (B) Real-time quantitative PCR was performed on RNA to detect SIRT2 gene expression in fly larvae of Da-Gal4>SIRT2 RNAi. c. western blot analysis of SIRT2 in lysate of different fly lines.

**Supplemental Figure 2.**
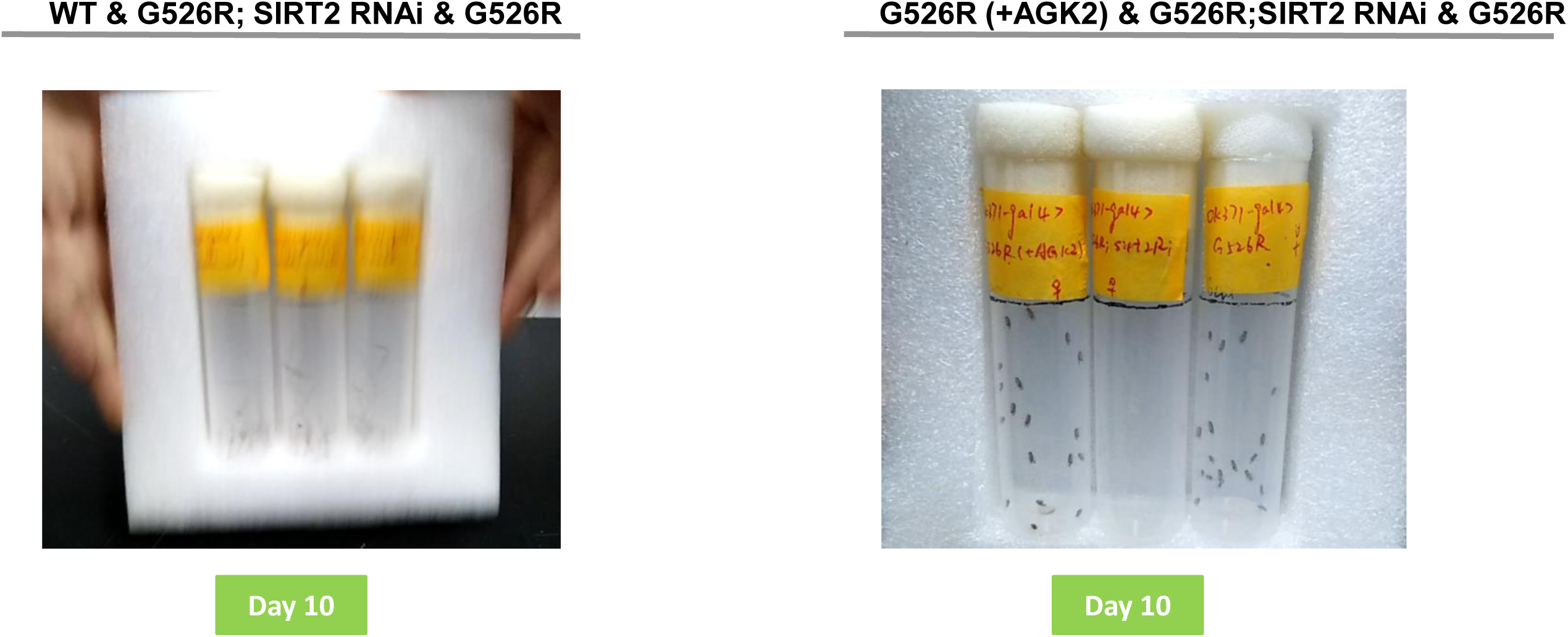
(A) Day 10 represents the motor performance analysis between the wildtype (left), SIRT2 RNAi in GARS^G526R^ and GARS^G526R^ flies. (B) Day 10 represents the motor performance analysis between the AGK2 treatment (left), SIRT2 RNAi in GARS^G526R^ and GARS^G526R^ flies. Bar graph displaying average climbing time to reach target in a negative geotaxis assay of female flies in motor neurons (OK371-GAL4). 4% DMSO was fed in all lines. N>100. Error bars represent s.e.m. ****P<0.0001.

